# Single-molecule imaging of IQGAP1 regulating actin filament dynamics in real time

**DOI:** 10.1101/2021.04.18.440338

**Authors:** Gregory J. Hoeprich, Shashank Shekhar, Bruce L. Goode

**Affiliations:** Department of Biology, Brandeis University, 415 South Street, Waltham, MA 02453, USA; Departments of Physics and Cell Biology, Emory University, Atlanta, GA 30322

**Keywords:** IQGAP1, actin, cytoskeleton, cell migration, TIRF microscopy, single molecule analysis

## Abstract

IQGAP is a conserved family of actin-binding proteins with essential roles in cell motility, cytokinesis, and cell adhesion, yet it has remained poorly understood how IQGAP proteins directly regulate actin filament dynamics. To close this gap, we used single-molecule and single-filament TIRF microscopy to directly visualize IQGAP regulating actin dynamics in real time. To our knowledge, this is the first study to do so. Our results show that full-length human IQGAP1 forms dimers that stably bind to filament sides and transiently cap barbed ends. These interactions organize actin filaments into thin bundles, suppress barbed end growth, and inhibit filament disassembly. Surprisingly, each activity depends on distinct combinations of IQGAP1 domains and/or dimerization, suggesting that different mechanisms underlie each functional effect on actin. These observations have important implications for how IQGAP functions as a direct actin regulator in vivo, and how it is deployed and regulated in different biological settings.

## INTRODUCTION

IQGAP is a large multi-domain actin-binding protein that is conserved across the animal and fungal kingdoms^1^, and plays crucial roles in cytokinesis, cell migration, phagocytosis, and cell adhesion^2-5^. The founding member of this protein family, human IQGAP1, was identified in 1994 and named based on its sequence similarity to GTPase-activating protein (GAP) proteins^6^. Subsequently, IQGAP1 was shown to interact with Cdc42 and Rac1, but was found to lack GAP activity. Instead, IQGAP1 stabilized Cdc42 in its active GTP-bound form^5,7^. Mammals have three IQGAP genes (IQGAP1-3), with IQGAP1 being the best characterized^8^. IQGAP1 functions directly downstream of Cdc42 and Rac1 at the leading edge, and is required for polarized cell migration and proper lamellipodial protrusion dynamics^5,7^. Upregulated IQGAP1 expression promotes motility^2^, and is associated with aggressive cancers and tumorigenesis^9-11^.

IQGAP1 is often referred to as a ‘scaffold’ protein because it associates with a number of different cytoskeletal regulatory proteins, including N-WASP, adenomatous polyposis coli (APC), CLIP-170, CLASP, and the formin Dia1 (Fig. 1A)^12-15^. However, IQGAP1 also directly binds to actin filaments^16-18^. Thus, a key step in understanding how IQGAP1 functions in vivo is to precisely define the kinetics of its interactions with actin filaments and its direct regulatory effects on actin filament dynamics. To date, only a single study has investigated the in vitro effects of an IQGAP protein on actin filament dynamics, using bulk pyrene-actin assembly assays to reveal that IQGAP1 slows barbed end growth and stabilizes filaments^16^. However, this has left open many questions about IQGAP’s activities and mechanism, which can be difficult to answer using bulk assays due to their inherent limitations. Bulk assays that monitor actin assembly kinetics fail to distinguish between effects on filament nucleation versus elongation, and bulk assays that monitor F-actin disassembly kinetics fail to distinguish between effects from severing versus depolymerization. Further, because bulk assays provide a readout of the average behavior of the entire filament population, they are unable to resolve two or more simultaneous effects on actin.

**Figure 1.**
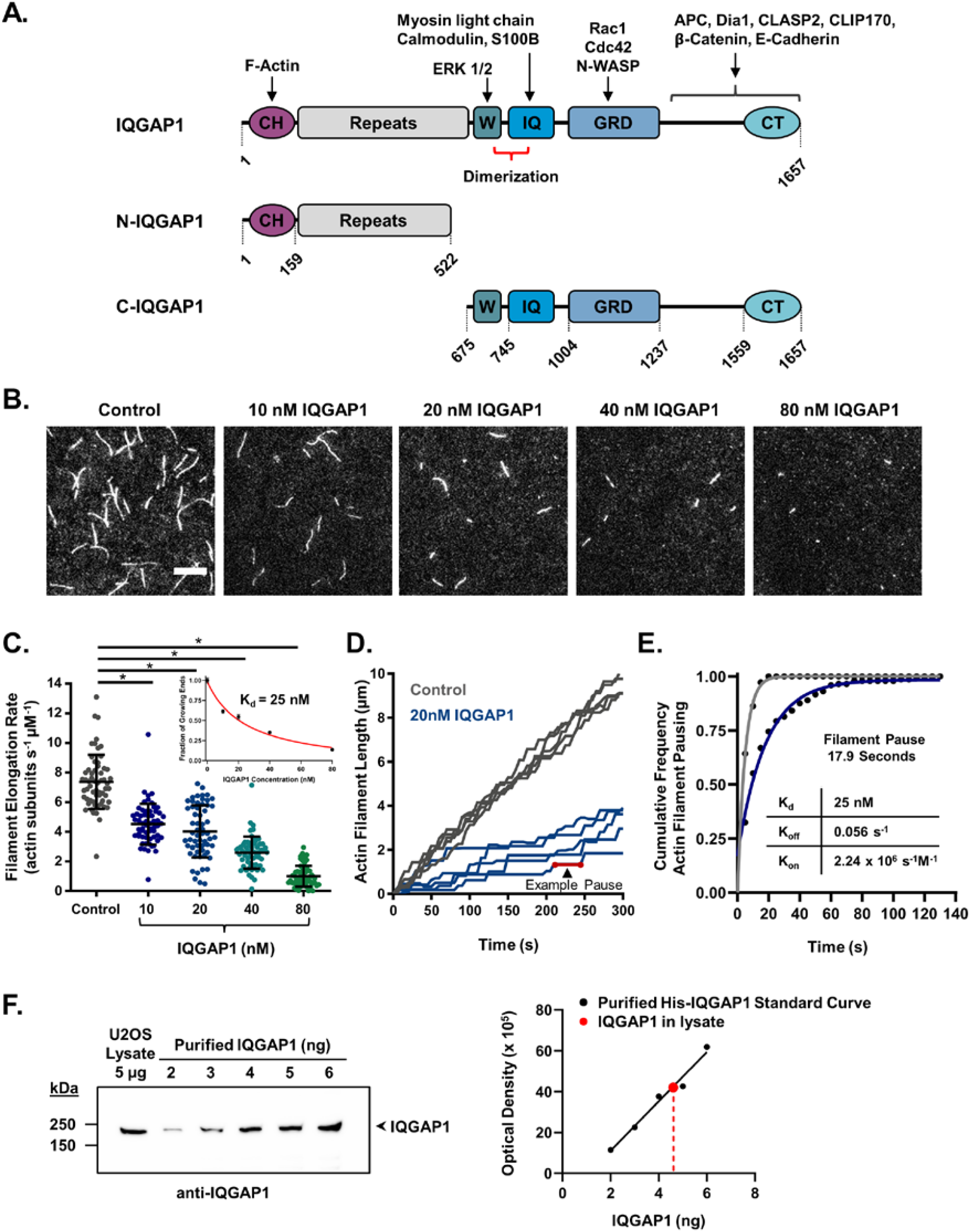
IQGAP1 transiently caps actin filament ends to inhibit barbed end growth. (**A**) Domain layouts for full-length IQGAP1 and fragments used in this study. Domains: CH, calponin homology; Repeats, six 50 amino acid repeats; W, WW domain; IQ, four isoleucine-glutamine motifs; GRD, GAP-related domain; CT, C-terminal domain. Amino acid numbering and boundaries are from UniProt: P46940, PDB: 1X0H and structural work^23,24^. Binding partners of different domains are shown. (**B**) Representative images from open-flow TIRF microscopy assays 10 min after initiation of actin assembly. Reactions contain 1 µM G-actin (10% Oregon green-labeled, 0.5% biotin-labeled) and different concentrations of full-length IQGAP1. Scale bar, 10 μm. (**C**) Barbed end elongation rates for actin filaments in TIRF reactions as in B (n=60 filaments, pooled from three independent trials for each condition). Mean and SD. Student’s t-test used to determine statistical significance of differences between conditions (* p < 0.05). Inset graph: fraction of free growing barbed ends versus concentration of IQGAP1 (nM) fit with a hyperbolic binding curve to measure equilibrium binding constant (K_d_=25 nM). Error bars, SEM. (**D**) Example traces of individual filament lengths over time (5 each) for control reactions and reactions containing 20 nM IQGAP1, from the same reactions as in C. Note the increase in pause time (no growth) in the presence of 20 nM IQGAP1 (example shown by red line). (**E**) Duration of pauses in the presence of 20 nM IQGAP1 (blue curve, n=38) compared to control (gray curve, n=176). The average barbed end pause time in the control reactions (lacking IQGAP1) was 4.7 s and in the presence of 20 nM IQGAP1 was 17.9 s. Fits were calculated from a single exponential equeation. Inset: table listing IQGAP1 binding affinity, on-rate, and off-rate for the barbed end. (**F**) Representative quantitative western blot (one of three independent trials) used to determine the concentration of endogenous IQGAP1 in U2OS cells. Blot was probed with anti-IQGAP1 antibody to compare the signal for endogenous IQGAP1 in the cell lysate lane to known quantities of purified 6His-IQGAP1. A standard curve was generated from the signals on the blot. The average cellular concentration of IQGAP1 (405 nM +/-112) was calculated from values obtained in three independent trials (482, 276, and 458 nM).

Here, we have overcome these limitations by using total internal reflection fluorescence (TIRF) microscopy to directly observe the effects of human IQGAP1 on the dynamics of individual actin filaments, and to observe single molecules of IQGAP1 interacting with filaments in real time^19,20^. Our results show that full-length IQGAP1 forms dimers that tightly associate with actin filament sides and: (i) transiently cap barbed ends to pause filament growth, (ii) organize filaments into thin bundles, and (iii) stabilize filaments against depolymerization. Further, we assign roles for the N-and C-terminal actin-binding halves of IQGAP1 in these activities, and provide additional evidence for distinct mechanisms underlying each regulatory effect on actin. Overall, our results provide new mechanistic insights into how IQGAP family proteins directly associate with and control actin filament dynamics and spatial organization, with important implications for IQGAP in vivo functions and regulation.

## RESULTS

### IQGAP1 transiently caps barbed ends of actin filaments to attenuate growth

We initiated our investigation by directly observing the effects of purified human IQGAP1 on actin filament barbed end growth using conventional open-flow TIRF assays. Oregon-green (OG) labeled actin filaments were polymerized in open-flow TIRF chambers, and sparsely tethered by incorporation of a low percentage of biotin-actin subunits (Fig. 1B). Monitoring polymerization allowed us to identify the fast-growing barbed ends and measure their rate of growth. In control reactions, barbed ends grew at 7.4 ± 1.8 subunits s^-1^ μM^-1^ (Fig. 1C), consistent with previous studies^21,22^. Addition of full-length IQGAP1 led to fewer and shorter filaments in the fields of view (Fig. 1B, Movie 1), and a concentration-dependent reduction in barbed end growth rate (Fig. 1C). These effects were potent, as 80 nM IQGAP1 was sufficient to strongly inhibit elongation (1.0 ± 0.7 subunit s^-1^ μM^-1^).

To better understand the mechanism of filament growth inhibition, we generated traces of filament length over time, focusing on reactions containing 20 nM IQGAP1, which exhibited an intermediate level of inhibition (Fig. 1D). Our reasoning was that these reactions would give us the best chance of detecting potential pauses in growth (capping events). This analysis revealed alternating phases of growth and no growth at filament barbed ends, suggesting that IQGAP1 transiently blocks barbed end growth, rather than persistently slowing growth. Consistent with this hypothesis, the barbed end growth rate that occurred between pauses was the same as the growth rate throughout control reactions lacking IQGAP1 (Fig. S1). Direct observation of filaments in real time was essential to uncovering the transient capping activity.

To determine the off-rate of IQGAP1 from barbed ends, we measured the durations of the pauses in growth induced by IQGAP1 (Fig. 1D). This yielded an off-rate of 0.056 s^-1^, corresponding to an average dwell time of approximately 18 seconds (Fig. 1E). By plotting the fraction of free growing ends versus IQGAP1 concentration, we also determined the equilibrium constant (K_d_) for IQGAP1’s association with barbed ends to be K_d_ = 25 nM (inset, Fig. 1C). Using the experimentally determined K_d_ and off-rate, we estimated the on-rate to be 2.24 × 10^6^ s^-1^ M^-1^ (Fig. 1E).

In order to determine whether the concentrations of IQGAP1 required to inhibit barbed end growth in vitro are physiologically relevant, we used quantitative western blotting to define the concentration of IQGAP1 in U2OS osteosarcoma cells (Fig. 1F). The average from three experiments was 405 nM +/-112 (mean and SD), which is well above the in vitro concentrations that strongly inhibited barbed end growth in our TIRF experiments. These values were also similar to the reported IQGAP1 concentration in MTD-1A epithelial cells (∼300 nM)^17^. Importantly, F-actin levels in mammalian cell lines are estimated to be >200 µM ^25,26^, suggesting that only a small percentage of the F-actin in cells could be decorated by IQGAP1, and this is consistent with the specificity of IQGAP1 localization to actin networks at the leading edge^5,7,18^. Further, IQGAP1 is sufficiently high enough in concentration in cells to efficiently cap barbed ends where it localizes, and we suggest that this capping activity may help promote the assembly of actin networks (see Discussion).

### Full-length IQGAP1 and its N-terminal half tightly bind to actin filament sides

To define the kinetics of IQGAP1 interactions with actin filaments, we purified and fluorescently-labeled SNAP-tagged full-length IQGAP1 (649-SNAP-IQGAP1). Importantly, addition of the tag and the dye did not alter IQGAP1 suppression of barbed end growth (Fig. S2A). We first analyzed the oligomeric state of our protein. Previous structural studies have suggested the presence of a strong dimerization activity in the W-IQ region (763-863) and a weaker dimerization activity in the N-terminus adjacent to the CH domain^17,24,27,28^. The presence of multiple dimerization domains in IQGAP1 has raised the possibility that the full-length protein might form higher-order oligomerization states beyond dimers. On the other hand, equilibrium sedimentation analysis has indicated that full-length IQGAP1 forms dimers^18^. As an independent test, we performed step-photobleaching analysis on purified 649-SNAP-IQGAP1 (with or without an N-terminal GST-tag) (Fig. 2 A & B, Fig. S2B). Our results show that full-length IQGAP1 forms stable dimers, with little evidence of higher-order oligomerization, agreeing well with the sedimentation analysis study above.

**Figure 2.**
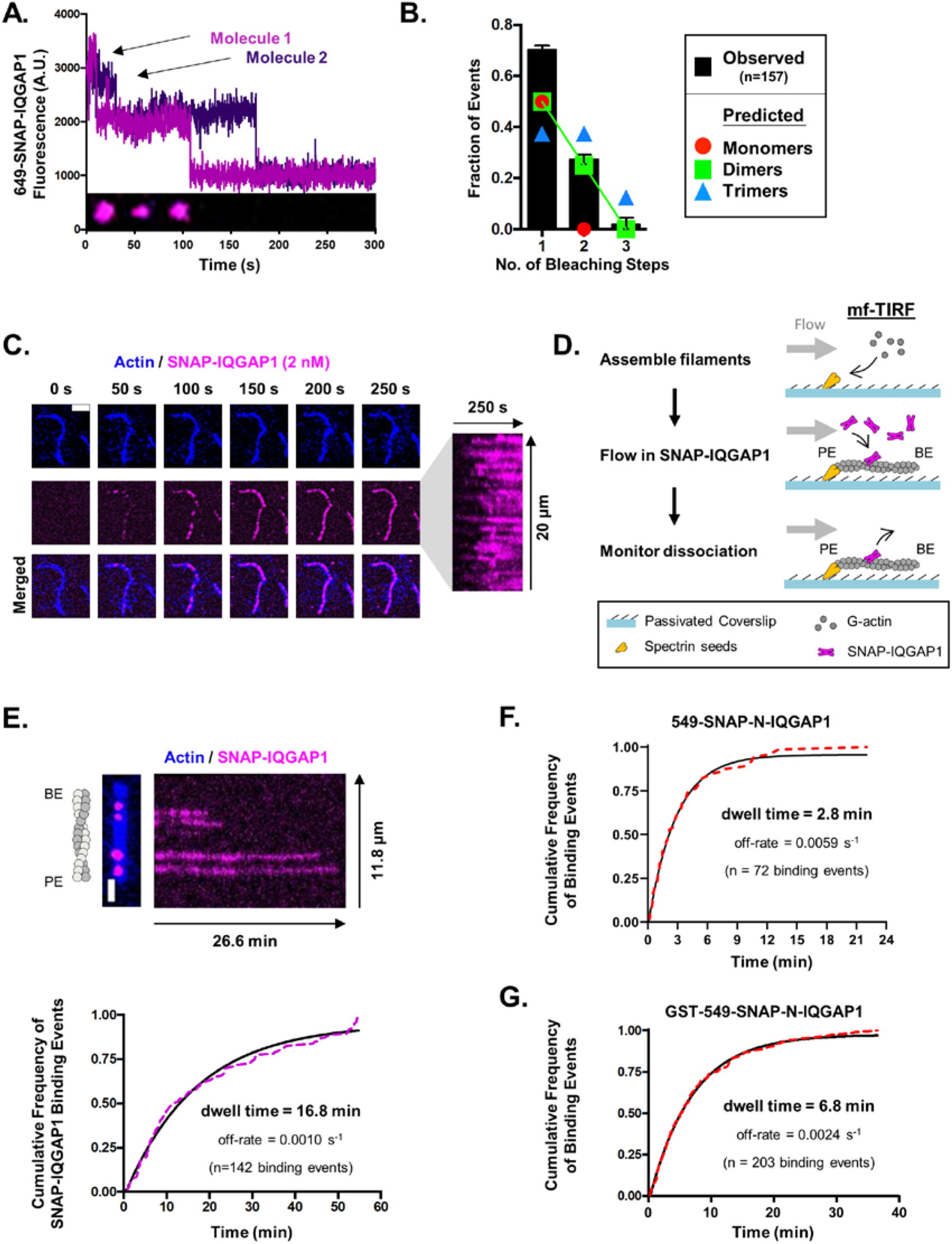
IQGAP1 dimers bind stably to the sides of actin filaments. (**A**) Representative step photobleaching traces from single molecules of full-length 649-SNAP-IQGAP1. Plot shows fluorescence intensity over time. Inset shows montage of images for one of the molecules shown in the plot (molecule 1, magenta). (**B**) Fraction of 649-SNAP-IQGAP1 molecules (n=157) that photobleached in one, two, or three steps (>3 photobleaching steps was never observed) from analysis as in A. Error bars, SEM. Observed fraction of photobleaching events (black) is compared to predicted fraction of photobleaching events (based on SNAP-labeling efficiency^21^) for different oligomeric states (colored coated symbols). (**C**) Representative time-lapse images and kymograph from TIRF reaction containing 2 nM 649-SNAP-IQGAP1, showing molecules (magenta) binding to an actin filament (blue). Scale bar, 2 μm. Kymograph shows 649-SNAP-IQGAP1 decoration is distributed along the filament over time. (**D**) Schematic showing experimental strategy to monitor 649-SNAP-IQGAP1 dissociation from filaments by microfluidic-assisted TIRF (mf-TIRF). Actin filaments with free barbed ends were polymerized from coverslip-anchored spectrin-actin seeds in the presence of 1 μM G-actin (15% Alexa-488-labeled) and 5 μM profilin, then capped at their barbed ends by flowing in 100 nM mouse capping protein (CP) for 1 min to prevent subsequent disassembly. Next, 0.5 nM 649-SNAP-IQGAP1 (without actin) was flowed in for 1 min to allow binding to filament sides, then buffer was flowed in (to remove free 649-SNAP-IQGAP1), and dissociation of 649-SNAP-IQGAP1 molecules was monitored over time. PE, pointed end; BE, barbed end. (**E**) Representative image and kymograph of 649-SNAP-IQGAP1 molecules (magenta) bound to an actin filament (blue). Scale bar, 2 μm. Observed dwell times (n=142 binding events) were plotted (dotted line), and an exponential fit (black line) was used to calculate the average dwell time of 16.8 min. (**F**) Observed dwell times of 549-SNAP-N-IQGAP1 molecules (n=72 binding events) were plotted (dotted line), and an exponential fit (black line) was used to calculate the average dwell time of 2.8 min. (**G**) Observed dwell times GST-549-SNAP-N-IQGAP1 molecules (n=203 binding events) were plotted (dotted line), and an exponential fit (black line) was used to calculate the average dwell time of 6.8 min.

We attempted to monitor 649-SNAP-IQGAP1 molecules interacting with filament sides using open-flow TIRF microscopy, where filaments were first assembled and tethered, and then a low concentration (2 nM) of 649-SNAP-IQGAP1 was flowed in. Under these conditions, we could readily detect binding of 649-SNAP-IQGAP1 molecules to filament sides early in the reactions. However binding was very stable, which meant that filaments steadily accumulated IQGAP1 on their sides, making it difficult to detect dissociation events (Fig. 2C). For this reason, we turned to using microfluidics-assisted TIRF (mf-TIRF), which allows new ingredients to be flowed in and out of the chambers, and aligns and straightens filaments under flow, providing more accurate measurements of filament length^29^. In these assays, we anchored filaments at their pointed ends to grow them by barbed end polymerization. We next briefly flowed in 649-SNAP-IQGAP1 (without actin monomers) to allow binding, and then washed free molecules out in order to monitor dissociation events in the absence of new binding events (Fig. 2D). The average lifetime of 649-SNAP-IQGAP1 binding on filament sides was 16.8 minutes, corresponding to an off-rate of 0.001 s^-1^ (Fig. 2E). To control for photobleaching effects, we repeated the analysis at a reduced frequency of image capture and obtained an off-rate that was not statistically different (Fig. S2C). Thus, full-length IQGAP1 interacts with actin filament sides very stably, indicative of a high affinity interaction.

To better understand which domains in IQGAP1 are responsible for its dimerization and interactions with actin filament sides, we purified and labeled a SNAP-tagged N-terminal fragment of IQGAP1 (1-522), with and without a GST tag (Fig. 1A). Based on previous studies, we expected that the non-GST-tagged N-IQGAP1 polypeptide would be monomeric, since it lacks the dimerizing W-IQ region^27^, and this was confirmed by step photobleaching analysis (Fig. S3A). Further, the GST-tagged 549-SNAP-N-IQGAP1 polypeptide was dimeric (Fig. S3B). Using mf-TIRF assays as above for full-length IQGAP1, we found that monomeric 549-SNAP-N-IQGAP1 molecules stably interacted with filament sides with an average dwell time of 2.8 minutes, corresponding to an off-rate of 0.006 s^-1^ (Fig. 2F). Dimeric GST-tagged 549-SNAP-N-IQGAP1 molecules had an average dwell time of 6.8 minutes, corresponding to an off-rate of 0.0025 s^-1^ (Fig. 2G). Together, our data show that the monomeric N-terminal half of IQGAP1 (1-522) is sufficient to stably bind actin filament sides, and suggests that the C-terminal half of IQGAP1 makes only a modest contribution to filament side-binding.

Finally, we also purified and labeled a SNAP-tagged C-terminal fragment of IQGAP1 (675-1657), 649-SNAP-C-IQGAP1. However, it did not bind to filament sides (Fig. S4A), and it failed to suppress barbed end growth (Fig. S4B), suggesting that the SNAP-tag likely interferes with its interactions with actin.

### Inhibition of barbed end growth by the N- and C-terminal halves of IQGAP1

Next, we asked whether the inhibitory effects of IQGAP1 on barbed end growth are mediated by its N-and/or C-terminal halves. Using open-flow TIRF microscopy, we compared the activities of different concentrations of N-IQGAP1 and C-IQGAP1 on the rate of growth at barbed ends (Fig. 3). Increasing concentrations of either fragment resulted in fewer and shorter filaments (Fig. 3A), and each fragment alone caused a concentration-dependent decrease in growth rate (Fig. 3B and 3C). Interestingly, the inhibitory effects of each fragment alone plateaued at ∼50% of the control rate of growth, whereas full-length IQGAP1 almost completely suppressed all growth (Fig. 3D). Furthermore, adding the two halves (N-IQGAP1 and C-IQGAP1) together *in trans* failed to improve the inhibitory effects beyond those of each fragment alone (Fig. 3C; also see arrow in Fig. 3D), suggesting that that full inhibition requires both halves of IQGAP1 physically linked in the same molecule.

**Figure 3.**
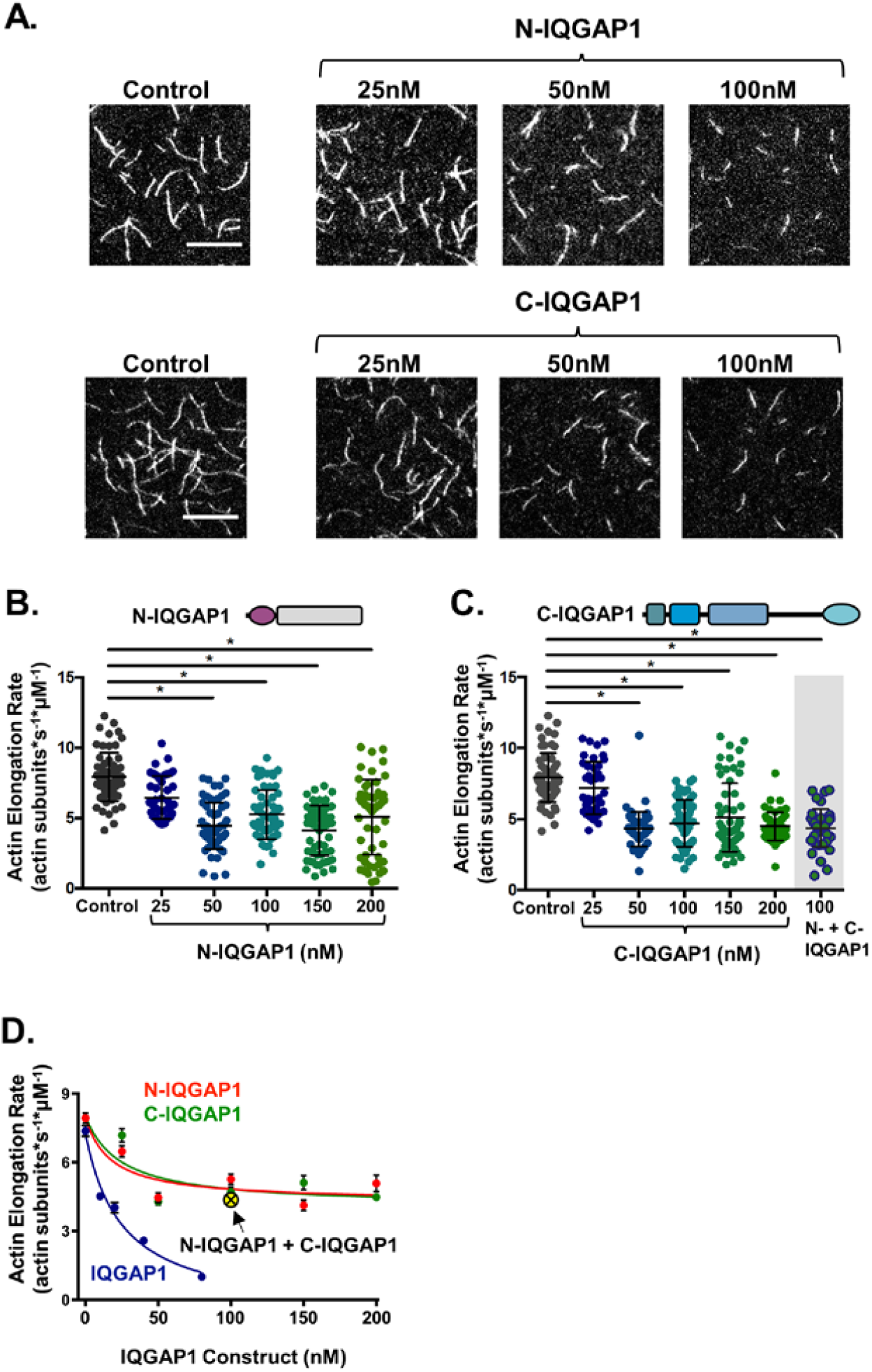
Each half of IQGAP1 partially suppresses actin filament growth. (**A**) Representative images from open-flow TIRF microscopy assays 10 min after initiation of actin assembly. Reactions contain 1 µM G-actin (10% Oregon green-labeled, 0.5% biotin-labeled) and different concentrations of N-IQGAP1 or C-IQGAP1. Scale bar, 10 μm. (**B**) Barbed end growth rates for filaments in TIRF reactions as in A, comparing the effects of different concentrations of N-IQGAP1. Data pooled from three independent trials (number of filaments analyzed for each condition, left to right: 60, 40, 60, 60, 60, 55). Mean and SD. Student’s t-test used to determine statistical significance of differences between conditions (* p < 0.05). **(C)** Same as B, except testing variable concentrations of C-IQGAP1 (number of filaments analyzed for each condition, left to right: 60, 40, 60, 60, 60, 55, 40). Gray shaded data show the combined effects of N-IQGAP1 and C-IQGAP1 (100 nM each) on barbed end elongation rate. (**D**) Comparison of concentration-dependent effects of full-length IQGAP1 (data from Figure 1C), N-IQGAP1 (data from B), and C-IQGAP1 (data from C) on barbed end growth rate. For each, the data were fit to single exponential decay curve. Error bars, SEM. Yellow dot highlights the combined effects of N-IQGAP1 and C-IQGAP1 (100 nM each).

### Dimerization of the N-terminal half of IQGAP1 promotes actin filament bundling

In addition, we used open-flow TIRF microscopy to examine how IQGAP1 affects the spatial organization of actin filaments. We first mixed labeled 649-SNAP-IQGAP1 with preassembled actin filaments (5-10 µm long) and observed that over time filament sides became increasingly decorated by IQGAP1 and grew thicker, i.e., formed bundles (Fig. 4A). To understand which domain(s) of IQGAP1 mediate bundling, we compared the effects of 10 nM full-length IQGAP1, N-IQGAP1, and C-IQGAP1 (Fig. 4B). N-IQGAP1 induced weak bundling compared to full-length IQGAP1, and C-IQGAP1 lacked significant bundling activity (Fig. 4B and 4C). The thickness of the bundles was assessed by two different methods: (i) measuring fluorescence intensity along the length of the bundles and calculating fluorescence density per micron of bundle length (Fig. 4C); (ii) measuring fluorescence intensity of a fixed-width line segment drawn perpendicular to the bundle (Fig. 4D). By each method, full-length IQGAP1 approximately tripled the fluorescence/thickness of filaments, suggesting formation of bundles approximately three filaments thick. In contrast, N-IQGAP1 only increased the fluorescence/thickness of filaments ∼1.5 fold, indicating a reduced bundling activity compared to full-length IQGAP1. Thus, bundling is substantially reduced in the absence of the C-terminal half of IQGAP1. We considered whether the C-terminus, which contains the dimerization domain, is important for bundling because it dimerizes the N-terminus. To test this idea, we compared the bundling activities of 10 nM monomeric N-IQGAP1 and GST-dimerized N-IQGAP1 (Fig. 4E). GST-tagged N-IQGAP1 organized filaments into bundles of similar thickness to those organized by full-length IQGAP1, suggesting that indeed dimerization is required for efficient bundling.

**Figure 4.**
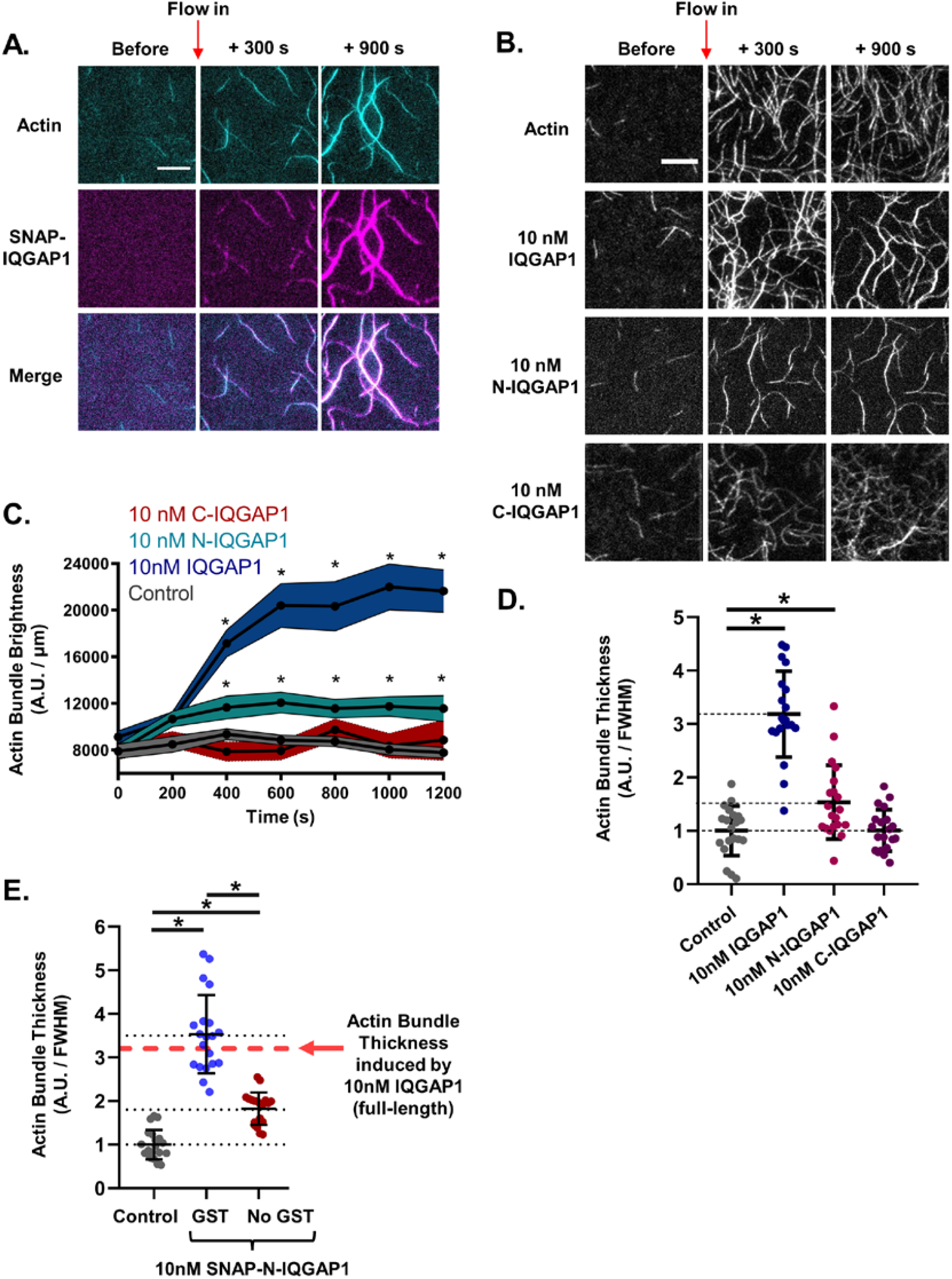
Dimerization of N-IQGAP1 promotes actin filament bundling. (**A**) Representative time-lapse images from open-flow TIRF microscopy reactions containing 2 μM F-actin (10% Oregon green-labeled) and 2 nM 649-SNAP-IQGAP1. Scale bar, 10 μm. (**B**) Representative time-lapse images from TIRF microscopy reactions containing 2 μM F-actin (10% Oregon green-labeled) and 10 nM IQGAP1, N-IQGAP1, or C-IQGAP1. Scale bar, 10 μm. IQGAP1 (or control buffer) was flowed-in 300 seconds after initiation of actin assembly, when filaments had grown to lengths of 5-10 μm. (**C**) Change in bundle thickness over time for reactions as in B, determined by measuring the fluorescence intensity along a bundle and calculating fluorescence density per unit length. Student’s t-test used to determine statistical significance of increase in fluorescence observed after time zero (* p < 0.05). (**D**)Bundle thickness was also assessed by measuring fluorescence intensity at full-width half-max of line segments drawn perpendicular to the bundle. The fluorescence intensity values were normalized to control (2 μM F-actin). Student’s t-test used to determine statistical significance of differences between conditions (* p < 0.05). (**E**) Comparing monomeric vs dimeric N-IQGAP1 fragments bundling actin filaments by measuring fluorescence intensity at full-width half-max of a line segments drawn perpendicular to the bundle. The fluorescence intensity values were normalized to control (2 μM F-actin). Student’s t-test used to determine statistical significance of differences between conditions (* p < 0.05).

### The monomeric N-terminal half of IQGAP1 stabilizes filaments against depolymerization

IQGAP1 has been shown to suppress dilution-induced actin filament disassembly in bulk assays^16^. Given that filaments depolymerize more rapidly from their barbed ends than pointed ends in the absence of actin monomers^30^, we postulated that IQGAP1 likely inhibits barbed end depolymerization. To directly test this model, we used mf-TIRF assays to monitor the effects of IQGAP1 on barbed end depolymerization. Filaments that were anchored at their pointed-ends were first polymerized, and then different concentrations of full-length IQGAP1 (without actin monomers) were flowed in, and depolymerization at the barbed end was monitored over time (Fig. 5A and B). At 1 nM IQGAP1, the depolymerization rate was reduced to only ∼10% of control rate (0.6 subunits s^-1^ ± 0.9 versus 6.0 subunits s^-1^ ± 2.9), and in the presence of 10 nM IQGAP1 depolymerization was almost undetectable (0.1 subunits s^-1^ ± 0.1) (Fig. 5B). The concentration of full-length IQGAP1 required for half-maximal change in depolymerization rate (IC_50_) was 0.1 nM (Fig. 5C).

**Figure 5.**
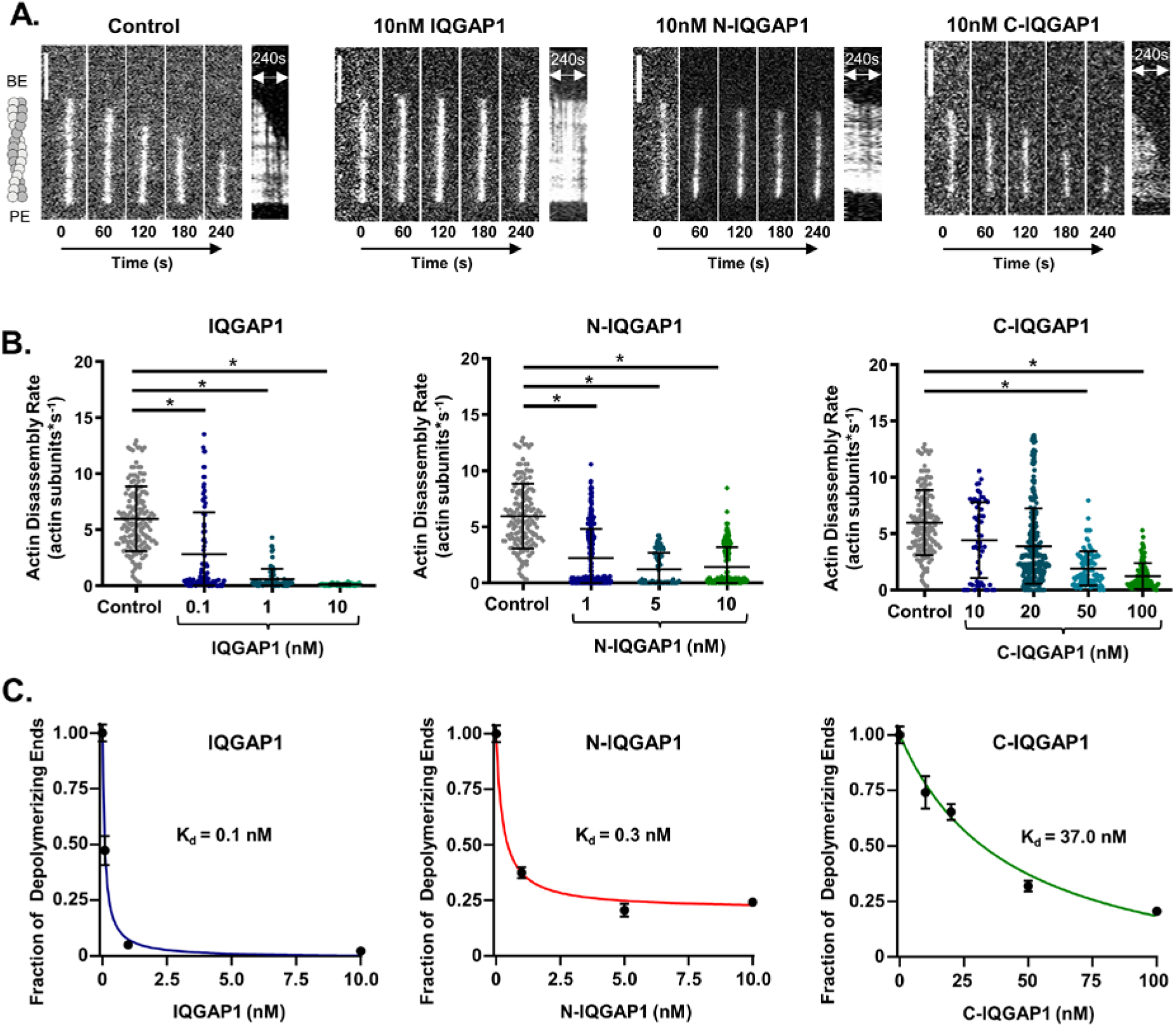
The monomeric N-terminal half of IQGAP1 strongly suppresses depolymerization at barbed ends. (**A**) Representative time-lapse images and kymographs of fluorescently labeled actin filaments (10% Oregon green-labeled actin) in mf-TIRF reactions, comparing depolymerization from barbed ends in the presence of 10 nM IQGAP1, N-IQGAP1, C-IQGAP1, or control buffer. Filaments anchored at their pointed ends were polymerized, and then IQGAP1, N-IQGAP1, or C-IQGAP1 (without actin monomers) was flowed in at time zero and depolymerization was monitored over time. Scale bar, 5 μm. (**B**) Barbed end depolymerization rates measured in the presence of different concentrations of IQGAP1, N-IQGAP1, and C-IQGAP1. Data pooled from three independent trials (number of filaments analyzed for each condition, left to right: 160, 93, 75, 68, 160, 315, 74, 221, 160, 59, 249, 107, 103). Mean and SD. Student’s t-test used to determine statistical significance of differences between conditions (* p < 0.05). (**C**) Graphs showing fraction of free depolymerizing barbed ends versus concentration of IQGAP1, N-IQGAP1, or C-IQGAP1, in which a hyperbolic binding curve was fit to the data to determine the equilibrium binding constant (K_d_). Error bars, SEM. Note that N-IQGAP1 is nearly as potent as full-length IQGAP1 in suppressing depolymerization, whereas C-IQGAP1 is ∼300-fold weaker.

Further analysis by mf-TIRF revealed that N-IQGAP1 potently stabilizes filaments against depolymerization at their barbed ends nearly as well as full-length IQGAP1 (Fig. 5B and C). In contrast, C-IQGAP1 was > 100-fold less potent than full-length IQGAP1 in stabilizing filaments. These observations suggest that monomeric N-IQGAP1 plays the dominant role in stabilizing filaments. This observation was somewhat unexpected, given the importance of C-IQGAP1 in inhibiting filament growth at barbed ends, and suggests that these two regulatory effects (inhibition of barbed end growth, and stabilization of filaments against depolymerization) have distinct underlying mechanisms.

## DISCUSSION

In a wide range of organisms IQGAP family proteins perform critical roles in controlling cellular actin dynamics, and yet there have been few in vitro studies to date investigating the nature of IQGAP’s direct interactions with actin and regulatory effects on actin filament dynamics. Our analysis using TIRF microscopy helps close this gap by providing what is to our knowledge the first direct visualization of an IQGAP family protein interacting with actin filaments, regulating their dynamics and spatial organization in real time. Below we discuss each of our mechanistic findings and their relevance to understanding of IQGAP’s in vivo roles as a direct actin regulator.

### IQGAP1 kinetic interactions with actin filaments

Previous studies using co-sedimentation assays hinted that IQGAP1 binds to actin filaments with high affinity, but did not quantify the interaction^18^. Using single molecule analysis, we directly observed and quantified the interactions of full-length human IQGAP1 molecules with the sides of filaments (dwell time

∼17 min; off-rate of 0.001 s^-1^). Given that full-length IQGAP1 dimerizes, we considered the possibility that its high affinity binding might stem from having two separate CH domains, since dissociation of IQGAP1 would then require simultaneous unbinding of both CH domains. However, we discovered that monomeric N-IQGAP1 is sufficient to tightly bind filament sides (dwell time ∼2.8 min; and off-rate of 0.006 s^-1^). Thus, our results suggest that a single CH domain may be sufficient for stable interactions with filament sides.

One in vivo implication of IQGAP family proteins having such a high affinity for actin filaments is that they may competitively block binding of other CH domain-containing proteins, e.g., fimbrin, filamin, calponin/transgelin, α-actinin, and MICAL^31^. Indeed, the cytokinetic actin ring (CAR) is strongly decorated by Rng2, the fission yeast homolog of IQGAP, and less-so by α-actinin and fimbrin^32^. The CH domain is also crucial for the essential function of *S. cerevisiae* IQGAP in cytokinesis^33^, consistent with its importance in directly binding actin filaments. The high affinity actin-binding interactions of IQGAP proteins further suggests that rapid reversal of their associations with actin networks in vivo may require negative control, e.g., through post-translational modifications modulating actin affinity^34^ and/or allosteric inhibition by ligands such as calmodulin^35,36^.

### Inhibition of barbed end growth

Using TIRF microscopy, we directly observed IQGAP1 inhibiting filament growth at barbed ends. Further, by analyzing the change in filament length over time we determined that IQGAP1 transiently caps barbed ends (dwell time ∼18 sec; off-rate 0.056 s^-1^; K_d_ = 25 nM). For comparison, conventional capping protein associates with barbed ends for tens of minutes^37,38^. Thus, IQGAP1 appears to be a transient capper.

Earlier bulk studies concluded that IQGAP1’s inhibitory effects on filament growth are mediated primarily by the C-terminal half of the protein^16^. However, we found that full inhibition of growth requires both halves of IQGAP1. Whereas 80 nM full-length IQGAP1 almost completely blocked barbed end growth, 200 nM of either half alone (N-IQGAP1 and C-IQGAP1) resulted in only ∼50% inhibition. Thus, our analysis reveals an important role for the N-terminal half in facilitating inhibition of barbed end growth. We considered whether the C-terminal half, which contains the dimerization domain (763-863)^27^, enhances capping simply by dimerizing monomeric N-IQGAP1. However, C-IQGAP1 alone was sufficient to inhibit barbed end growth equally well to N-IQGAP1. Thus, N-IQGAP1 and C-IQGAP1 make separate and independent contributions to the inhibition of filament growth. Future structural studies will be required to determine the underlying mechanisms. However, it is possible that binding of the CH domain of N-IQGAP1 to actin filament sides slows addition of new subunits via allosteric effects. Indeed, Hayakawa and co-workers have reported that binding of the N-terminus of Rng2, the *S. pom*be homolog of IQGAP, alters the structure of actin filaments^39^. Although the actin-binding domain of C-IQGAP1 is not as well characterized as the CH domain, it is required for full inhibition of growth. Further, it must be linked to N-IQGAP1 in the same molecule in order to do so (Fig. 3D), suggesting that coordination between the two actin-binding domains of IQGAP1 is required for this activity (see model, Fig. 6).

**Figure 6.**
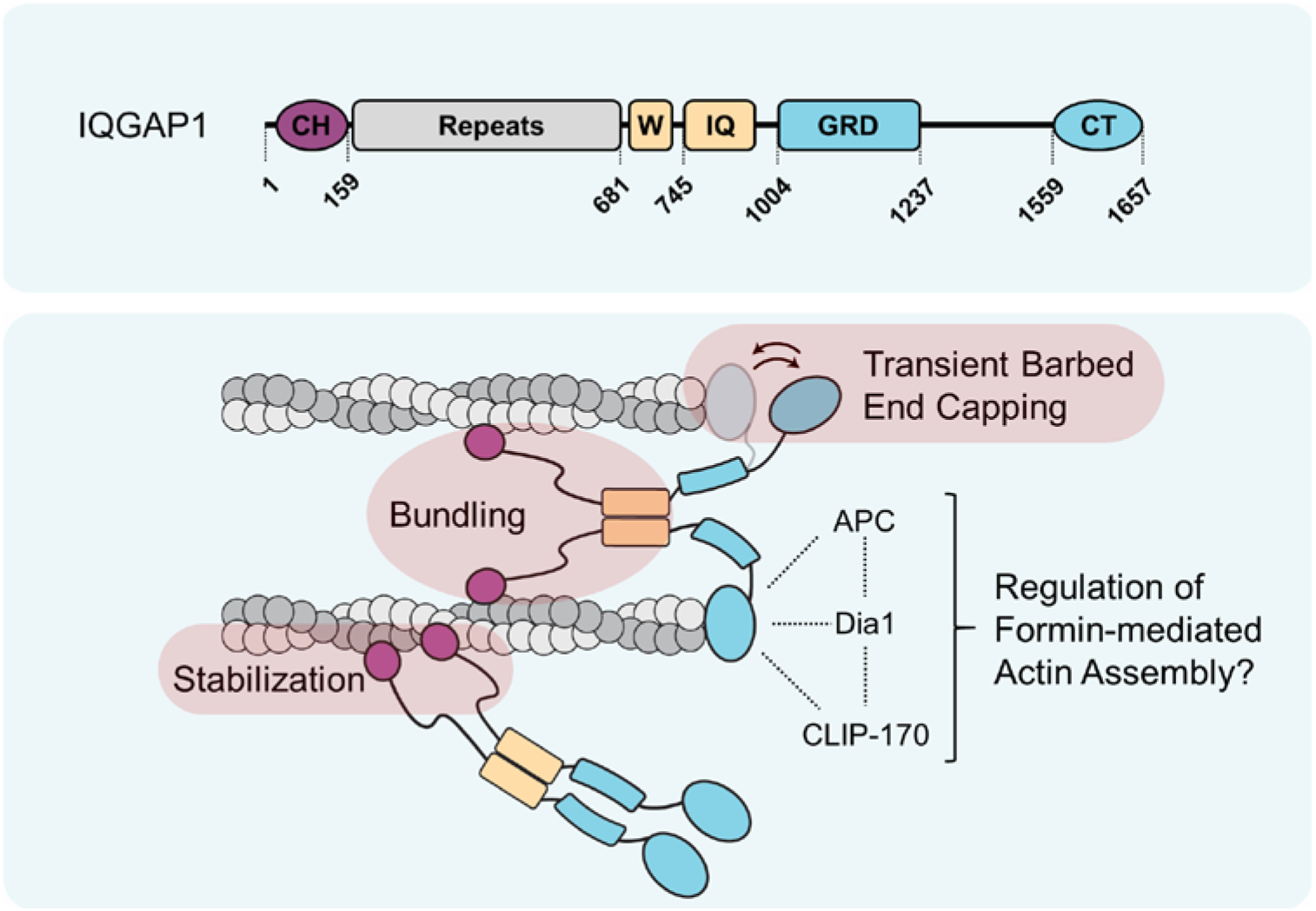
Working model for IQGAP1 regulatory activities on actin filament dynamics and spatial organization. Top panel shows domain layout of full-length IQGAP1. Bottom panel shows working model for how IQGAP1 dimers directly control actin filament growth, bundling, and stabilization, with each activity highlighted in red. The N-terminal half of IQGAP1 binds tightly to actin filament sides using its CH domain and plays a central role in stabilizing filaments. Dimerization of the N-terminal half is mediated by the W-IQ region of IQGAP1, which is required for bundling but not stabilization. C-terminal domains in IQGAP1 transiently cap barbed ends to facilitate inhibition of filament growth, but work in close coordination with N-terminal side-binding domains of IQGAP1 to achieve full inhibition. The C-terminal (CT) domain of IQGAP1 binds to the formin Dia1, as well as CLIP-170 and adenomatous polyposis coli (APC), which directly collaborate with Dia1 to promote actin assembly^12,13,15,21,47-49^. Thus, IQGAP1 may have additional regulatory roles in controlling formin-and APC-mediated actin assembly.

How might IQGAP1’s transient capping activity contribute its in vivo functions? IQGAP1 accumulates at the leading edge of cells and is required for normal lamellipodia protrusion velocity and frequency^2,5,7,18^. Furthermore, IQGAP1 is thought to promote actin assembly at the leading edge through interactions with N-WASP^14,40^ and Dia1^12^. How might transient capping by IQGAP1 contribute this role in promoting actin network assembly? Importantly, while capping suppresses the growth of individual actin filaments in a purified system, in the cellular context it is not synonymous with ‘negative regulation’. This is because capping in vivo focuses actin monomer addition to the newly-nucleated barbed ends via a ‘funneling’ effect^41,42^, and elevates actin monomer concentrations to help promote nucleation^43^. Thus, under cellular conditions capping activities can be instrumental in promoting actin network assembly.

### Actin filament bundling

Earlier studies using electron microscopy and falling ball viscosity assays demonstrated that IQGAP1 crosslinks actin filaments, and that this activity is mediated by N-terminal CH domain-containing fragments of the protein (1-216)^17^. Fukata and co-workers found that the minimal construct (1-863) that crosslinks filaments included both the CH domain and the suggested dimerization domain (763-863)^17,27^. In our TIRF assays, we observed IQGAP1 potently bundling actin filaments in real time (10 nM was sufficient to bundle 2 µM F-actin). Further, we measured bundle thickness and found that full-length IQGAP1 organizes filaments into thin bundles only a few filaments thick. In agreement with Fukata et al., we found that bundling requires dimerization of the N-terminal half of IQGAP1, either by a GST tag or inclusion of the C-terminal half which contains the dimerization domain (see model, Fig. 6). Bundling by IQGAP1 may be important for its role in promoting cell motility, cell adhesion, and cytokinesis. Further, bundling by IQGAP1 may be regulated in vivo, and indeed calmodulin binding inhibits IQGAP1’s bundling effects^36^. Further, Cdc42 and Rac1 binding to the GRD region of IQGAP1 may lead to its higher-order oligomerization^17^, potentially expanding or transforming its filament crosslinking capabilities.

### Stabilization of actin filaments

Using TIRF, we observed IQGAP1 potently stabilizing actin filaments against depolymerization in a concentration-dependent manner (Fig. 5B). These results validate earlier observations from bulk studies^16^, but extend our understanding of the mechanism. Many CH domain-containing proteins crosslink filaments as well as stabilize them against depolymerization, e.g., calponin, fimbrin, and IQGAP1^16,44^. This has suggested that crosslinking and stabilization activities may be coupled in CH domain family proteins. Here, we directly tested this model by monitoring the stabilization effects of IQGAP1 on single (non-bundled) actin filaments in mf-TIRF assays. This enabled us to uncouple stabilization and bundling, which is not possible to do using bulk assays, and our data show that IQGAP1 stabilizes filaments independent of bundling. Further, we found that monomeric N-IQGAP1 has a similar potency to full-length IQGAP1 in attenuating depolymerization (IC_50_ = 0.1 nM and 0.3 nM, respectively), yet has minimal bundling activity (Fig. 4E). Together, these observations suggest that IQGAP1 uses distinct mechanisms to promote filament stabilization and cross-linking, and that these effects can occur independently of one another (Fig. 6). This has important implications for IQGAP1 functions in vivo, as it suggests that these two activities could be independently regulated to differentially control actin network dynamics versus spatial organization.

### Concluding remarks

In this study, we have used direct visualization to define the kinetics of IQGAP1 interactions with actin filament sides and barbed ends, determined the contributions of each half of the protein to these interactions, and defined IQGAP1’s direct effects on actin filament dynamics. Our results show that IQGAP1 is a high affinity actin-binding protein with potent effects in stabilizing filaments and suppressing barbed end growth. As discussed above, these activities help to explain the previously demonstrated in vivo roles of IQGAP proteins in promoting actin assembly to facilitate such processes as cell migration, cell adhesion, and cytokinesis. Interestingly, another recent study performed in parallel to ours shows that IQGAP proteins tethered to lipid membranes generate actin filament structures that are highly curved, e.g., arcs and full rings^45^. Together, our studies provide an important framework for future investigations that address how IQGAP1 works together with in vivo binding partners to regulate actin dynamics at the leading edge, e.g., calmodulin, Cdc42, Dia1, APC, and CLIP-170^7,12-14,18,34,46^. Several of these ligands interact directly with the globular C-terminal domain of IQGAP1, suggesting that their activities may be coordinated with the transient barbed end capping activity of IQGAP1 to control actin assembly dynamics (Fig. 6).

## DATA AVAILABILITY

Data supporting the findings of this manuscript are available from the corresponding author upon reasonable request.

## ACKNOWLEGEMENTS

We are grateful to Johnson Chung for assistance with labeling proteins, Jeff Gelles for technical and intellectual input throughout the project, and Colby Fees and Amy Sinclair for valuable discussions, comments on the manuscript, and assistance with data analysis. We thank Marie-France Carlier for generously providing IQGAP1 plasmids, and the Brandeis microfabrication facility for assistance with generating PDMS microfluidic chips for mf-TIRF experiments. This work was supported by an NIH R35 award (GM134895) to B.L.G., and by the Brandeis NSF Materials Research Science and Engineering Center grant (DMR-1420382).

## AUTHER CONTRIBUTIONS

GJH and BLG designed the experiments and wrote the manuscript. GJH and SS performed experiments and analyzed the data.

## SUPPLEMENTAL MATERIALS

**Figure S1.**
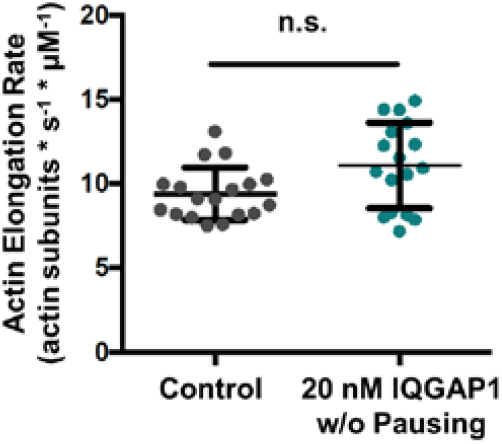
Additional controls supporting a role for IQGAP1 in inhibiting barbed end growth. Actin filament elongation rates during growth phases (between pausing events) from open-flow TIRF reactions in the presence of 20 nM IQGAP1 compared to control reactions. Figure 1D shows example filament traces, highlighting alternating pauses and growth phases in the presence of 20 nM IQGAP1. After excluding pause times, filaments in both reactions elongate at a similar rate, supporting the view that IQGAP1 transiently caps barbed ends to pause growth. Data averaged from two independent trials (control, n=18 filaments; IQGAP1, n=17 filaments). Error bars, SD. Student’s t test used to determine no significant difference (n.s.) between the two rates.

**Figure S2.**
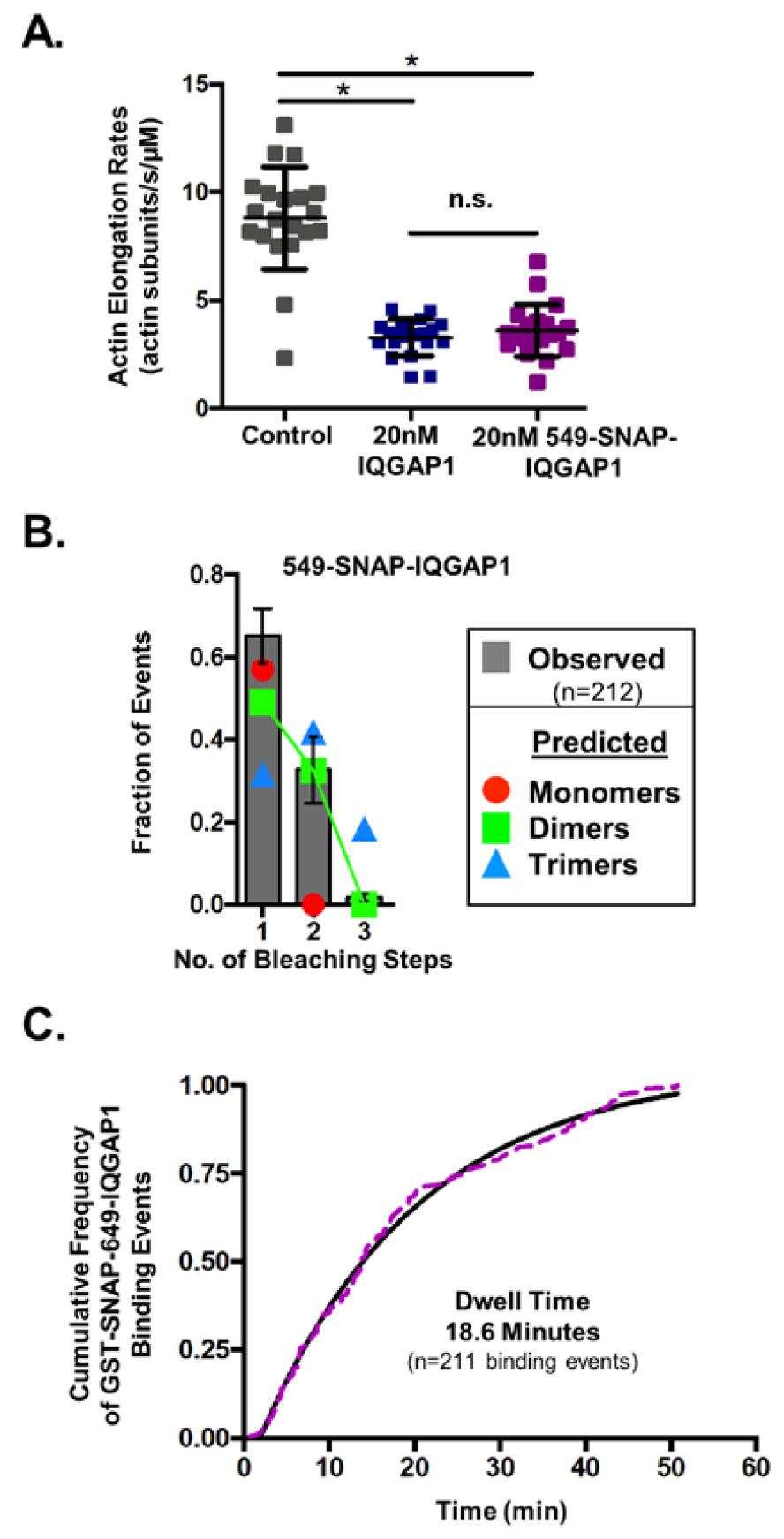
Controls for SNAP-tagged IQGAP1. (**A**) Effects of IQGAP1 versus 649-SNAP-IQGAP1 on rate of actin filament elongation in TIRF assays. Reactions contain 1 μM G-actin (10% Oregon green-labeled, 0.5% biotin-labeled) with 20 nM IQGAP1, 20 nM 649-SNAP-IQGAP1, or control buffer. Data averaged from two independent trials (n=20 for each condition). Error bars, SD. Student’s t-test used to determine statistical significance between conditions (* p < 0.05; n.s., not significant). (**B**) Removing the GST tag from SNAP-IQGAP1 does not alter its oligomerization state. Fraction of 549-SNAP-IQGAP1 molecules (n=212) that photobleached in one, two, or three steps (>3 photobleaching steps was never observed). Error bars, SEM. Observed fractions of photobleaching events (black) are compared to predicted fractions of photobleaching events (based on SNAP-labeling efficiency^21^) for different oligomeric states (colored coated symbols). Note: this SNAP-IQGAP construct, lacking a GST tag, forms dimers, similar to the GST-tagged SNAP-IQGAP1 construct (Figure 2B). (**C**) Dwell times of 649-SNAP-IQGAP1 molecules on the sides of actin filaments (n=211 binding events) measured as in figure 2E, except that images were acquired every 30 s instead of every 10 s, as a control for photobleaching effects. Data were plotted (dotted line), and an exponential fit (solid black line) was used to calculate the average dwell time of 18.6 min. A non-parametric Monte Carlo permutation resampling scheme was used to assess the statistical significance between the dwell times acquired at 30 s intervals (this figure) versus 10 s intervals (Figure 2E) ^50^, and no significant difference was found (p = 0.18).

**Figure S3.**
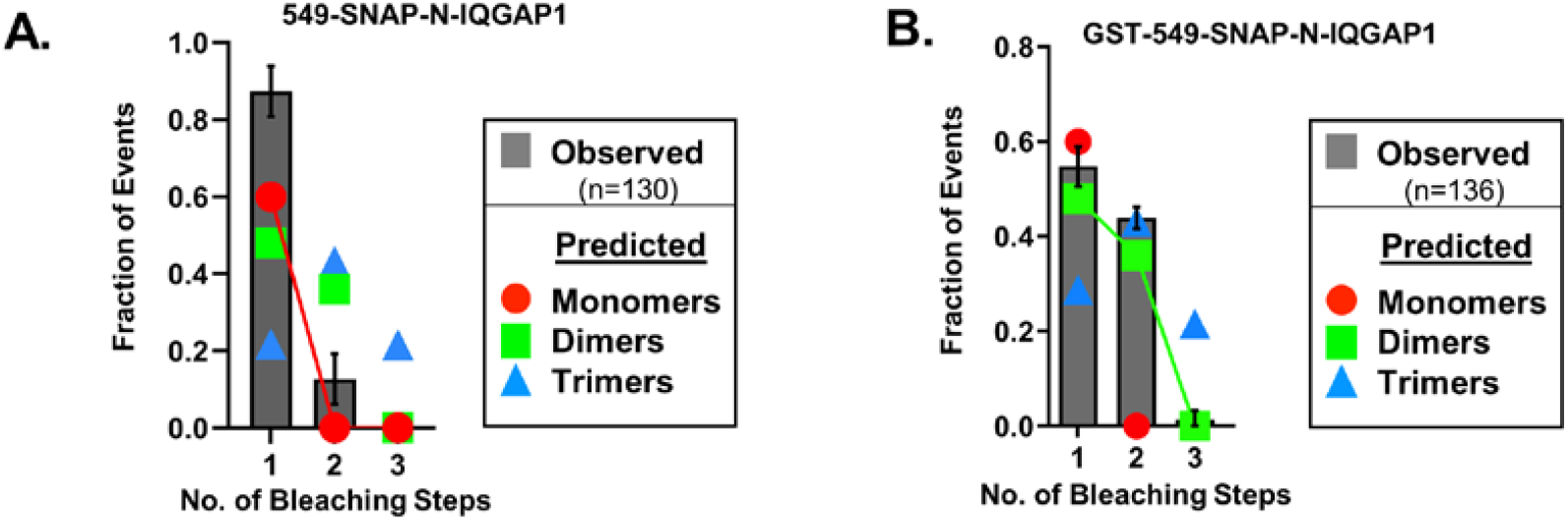
N-IQGAP1 is monomeric. (**A**) Step photobleaching analysis. Fraction of 549-SNAP-N-IQGAP1 molecules (n=136) that photobleached in one or two steps (>2 photobleaching steps was never observed). Error bars, SEM. Observed fractions of photobleaching events (grey) are compared to predicted fractions of photobleaching events (based on SNAP-labeling efficiency^21^) for different oligomeric states (colored coated symbols). (**B**) Fraction of GST-549-SNAP-N-IQGAP1 molecules (n=136) that photobleached in one, two, or three steps (>3 photobleaching steps was never observed). Error bars, SEM. Observed fractions of photobleaching events (grey) are compared to predicted fractions of photobleaching events (based on SNAP-labeling efficiency^21^) for different oligomeric states (colored coated symbols).

**Figure S4.**
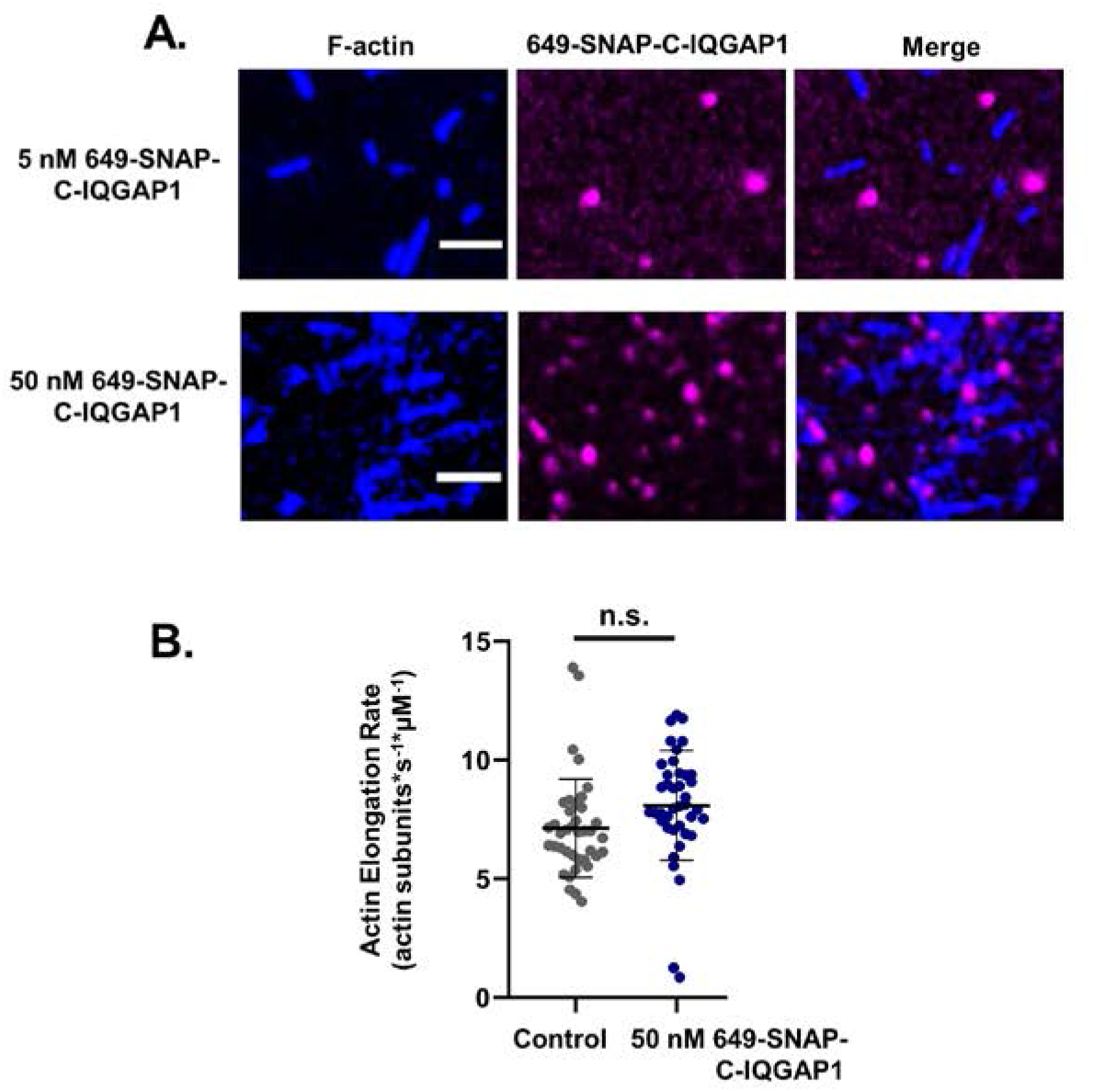
649-SNAP-C-IQGAP1 does not interact with actin. (**A**)Representative images from open-flow TIRF assays showing that 649-SNAP-C-IQGAP1 at two different concentrations (5 nM and 50 nM) does not visibly interact with actin filaments. Actin filaments were assembled and anchored using 1 μM G-actin (10% Oregon green-labeled, 0.5% biotin-labeled), and then 5 nM or 50 nM 649-SNAP-C-IQGAP1 was flowed in, and binding was monitored. Scale bar, 10 μm. (**B**) Effects of 649-SNAP-C-IQGAP1 on rate of actin filament elongation in TIRF assays. Reactions contain 1 μM G-actin (10% Oregon green-labeled, 0.5% biotin-labeled) with 50 nM 649-SNAP-C-IQGAP1 or control buffer. Data averaged from two independent trials (n=40 per condition). Error bars, SD. Student’s t-test used to determine statistical significance between conditions (n.s., not significant).

**Movie S1. Open-flow TIRF microscopy assays showing concentration-dependent effects of full-length IQGAP1 in inhibiting actin filament growth. Related to Figure 1**. Reactions contain 1 μM G-actin (10% Oregon green-labeled; 0.5% biotin-labeled) with different concentrations of full-length IQGAP1 as indicated. Video playback is 10 frames per second. Scale bar, 10 μm. Time stamp, min:sec.

**Movie S2. Open-flow TIRF microscopy assays showing 649-SNAP-IQGAP1 decorating and bundling actin filaments. Related to Figure 4**. Actin filaments were first assembled using 2 μM G-actin (10% Oregon green-labeled) until they had reached lengths of 5-10 μm, and then (∼200s into the movie) 2 nM 649-SNAP-IQGAP1 (magenta) was flowed in, and decoration and bundling of filaments (cyan) was monitored over time. Video playback is 10 frames per second. Scale bar, 10 μm. Time stamp, min:sec.

**Movie S3. Microfluidics-assisted TIRF (mf-TIRF) microscopy assays showing the effects of full-length IQGAP1 on the rate of filament depolymerization at barbed ends. Related to Figure 5**. Filaments anchored at their pointed ends with spectrin-actin seeds were polymerized at their free barbed ends using 1 μM G-actin (10% Alex488-labeled) until they were ∼10 μm long, and then 10 nM full-length IQGAP1 (without actin monomers) or control buffer was flowed in (at time zero in the movie) and depolymerization at barbed ends (yellow chevrons) were monitored over time. Video playback is 10 frames per second. Scale bar, 2 μm. Time stamp, min:sec.

## MATERIALS AND METHODS

### Plasmid construction

Plasmids for *E. coli* expression and purification of human full-length 6His-IQGAP1 (1-1657), 6His-N-IQGAP1 (1-522) and GST-C-IQGAP1 (675-1657) were generously provided by Dr. Marie-France Carlier (CNRS, Paris). The resulting tagged proteins are referred to as IQGAP1, N-IQGAP1, and C-IQGAP1 throughout this study; the GST tag (on C-IQGAP1) was removed only where specifically indicated. To generate plasmids for *E. coli* expression and purification of the same three IQGAP1 polypeptides with SNAP tags, coding regions from the plasmids above were PCR amplified and subcloned into the GST-pp-SNAP-pGEX-6p-1 vector^21^, which introduces an N-terminal GST tag, PreScission Protease site (pp), and SNAP tag, and a C-terminal 6His tag. SNAP-IQGAP1 proteins used in this study include all of these tags, except where it is noted that the GST tag was removed.

### Protein purification

Rabbit skeletal muscle actin (RMA) was purified from acetone powder^51^ generated from frozen ground hind leg muscle tissue of young rabbits (Pel-Freez Biologicals, Rogers, AR). Lyophilized acetone powder stored at −80°C was mechanically sheared in a coffee grinder, resuspended in G-buffer (5 mM Tris-HCl, pH 8.0, 0.2 mM ATP, 0.5 mM dithiothreitol (DTT), 0.1 mM CaCl_2_), and then cleared by centrifugation for 20 min at 50,000 × *g*, 4°C). The supernatant was filtered through Grade 1 Whatman paper, then the actin was polymerized by the addition of 2 mM MgCl_2_ and 50 mM NaCl to the filtrate and overnight incubation at 4°C with slow stirring. The next morning, NaCl powder was added to a final of 0.6 M, and stirring was continued for another 30 min at 4°C. F-actin was pelleted by centrifugation for 150 min at 120,000 × *g*, 4°C. The pellet was solubilized by dounce homogenization and dialyzed against 1 liter of G-buffer at 4°C (three consecutive times at 12-18 h intervals). Monomeric actin was then precleared for 30 min at 435,000 × *g*, 4°C, and loaded onto a S200 (16/60) gel-filtration column (GE Healthcare; Marlborough, MA). Peak fractions containing actin were stored at 4°C.

For preparing biotinylated actin used in open-flow cell TIRF microscopy assays, the F-actin pellet above was dounced and dialyzed against G-buffer lacking DTT. Monomeric actin was then polymerized by the addition of an equal volume of 2x labeling buffer (50 mM imidazole pH 7.5, 200 mM KCl, 0.3 mM ATP, and 4 mM MgCl_2_). After 5 min, the actin was mixed with a fivefold molar excess of NHS-XX-Biotin (Merck KGaA, Darmstadt, Germany) and incubated for 15 h at 4°C. The F-actin was pelleted as above, and the pellet was rinsed with G-buffer, then homogenized with a dounce, and dialyzed against G-buffer for 48 h at 4°C. Biotinylated monomeric actin was purified further on an S200 (16/60) gel-filtration column as above. Aliquots of biotin actin were snap frozen in liquid N_2_ and stored at −80°C.

For the fluorescently labeled actin used in open-flow cell TIRF microscopy assays, actin was labeled on cysteine 374 as previously described^52^. Briefly, the F-actin pellet described above was dounced and dialyzed against G-buffer lacking DTT. Monomeric actin was then polymerized by adding an equal volume of 2x labeling buffer (50 mM Imidazole pH 7.5, 200 mM KCl, 0.3 mM ATP, 4 mM MgCl_2_). After 5 min, the actin was mixed with a five-fold molar excess of Oregon green (OG)-488 iodoacetamide (Life Technologies; Carlsbad, CA) resuspended in anhydrous dimethylformamide, and incubated in the dark for 15 h at 4°C. Labeled F-actin was pelleted as above, and the pellet was rinsed briefly with G-buffer, then depolymerized by dounce homogenization, and dialyzed against G-buffer for 2 days at 4°C. Labeled, monomeric actin was purified further on a 16/60 S200 gel-filtration column as above. OG-488-actin was dialyzed for 15 h against G-buffer with 50% glycerol and stored at −20°C. The concentration and labelling efficiency was determined by measuring the absorbance at 280 nm and 496 nm, using these molar extinction coefficients: ε_280_ actin = 45,840 M^-1^ cm^-1^, ε_496_ OG-488 = 76,000 M^-1^ cm^-1^, and OG-488 correction factor at 280 = 0.12.

For the fluorescently labeled actin used in microfluidics-assisted TIRF (mf-TIRF) assays, actin was labeled on surface-exposed primary amines as previously described^29^. Briefly, G-actin was polymerized by dialyzing overnight against modified F-buffer (20 mM PIPES pH 6.9, 0.2 mM CaCl_2,_ 0.2 mM ATP, 100 mM KCl). Then the F-actin was incubated for 2 h at room temperature with a five-fold molar excess of Alexa-488 NHS ester dye (Life Technologies). F-actin was then pelleted by centrifugation at 450,000 × *g* for 40 min at room temperature. The pellet was resuspended in G-buffer, and homogenized with a dounce, and incubated on ice for 2 h to depolymerize filaments. Actin was then re-polymerized on ice for 1 h after adding KCl and MgCl_2_ (final concentration of 100 mM and 1 mM respectively). F-actin was pelleted by centrifugation for 40 min at 450,000 × *g* at 4°C. The pellet was homogenized with a dounce and dialyzed overnight at 4°C against 1 liter of G-buffer. Next, the solution was centrifuged for 40 min at 450,000 × *g* at 4°C, and the supernatant was collected. The concentration and labelling efficiency was determined by measuring the absorbance at 280 nm and 495 nm, using these molar extinction coefficients: ε_280_ actin = 45,840 M^-1^ cm^-1^, ε_495_ Alexa-488 = 71,000 M^-1^ cm^-1^ and ε_280_ AF488 = 7,810 M^-1^ cm^-1^.

Human Profilin-1 was expressed in *E. coli* BL21 DE3 by growing cells to log phase at 37°C in Terrific Broth (TB) media and inducing expression with 1 mM IPTG at 37°C for 3 h. Cells were harvested by centrifugation and pellets were stored at −80°C. Cell pellets were resuspended in lysis buffer (50 mM Tris-HCl, pH 8.0, 1 mM EDTA, 0.2% Triton X-100, lysozyme,1 mM PMSF, and protease inhibitor cocktail: 0.5 µM each of pepstatin A, antipain, leupeptin, aprotinin, and chymostatin), and kept on ice for 30 min. Lysates were cleared for 30 min at 272,000 × *g* at 4°C, and the supernatant was collected and fractionated on a HiTrap Q column (GE Healthcare; Chicago, IL) equilibrated in 20 mM Tris, pH 8.0, and 50 mM NaCl and eluted with a salt gradient (0–1 M NaCl and 20 mM Tris, pH 8.0). Peak fractions were concentrated and then purified further on a Superdex 75 column equilibrated in 20 mM Tris-HCl pH 8.0, and 50 mM NaCl. Peak fractions were pooled, snap frozen in aliquots, and stored at −80°C.

Mouse non-muscle capping protein (CPa1b2 or CP) was purified as described^53^. Briefly, the expression vector^54^ was expressed in *E. coli* BL21 pLysS by growing cells to log phase at 37°C in Lauryl Broth media and inducing expression with 0.4mM IPTG at 37°C for 3 h. Cells were harvested by centrifugation and pellets were stored at −80°C. Cell pellets were resuspended in lysis buffer (20 mM Tris pH 8.0, 1 mM EDTA, 0.1% Triton X-100, lysozyme, a standard mixture of protease inhibitors), and kept on ice for 30 min. Lysates were cleared for 30 min at 12,500 × *g* at 4°C and the supernatant was collected and fractionated on a 1 ml Q-HiTrap column (GE Healthcare) equilibrated in 20 mM Tris, pH 8.0, and 50 mM NaCl and eluted with a salt gradient (0–0.5 M NaCl and 20 mM Tris, pH 8.0). Peak fractions were concentrated and then purified further on a Superdex 75 gel filtration column (GE Healthcare) equilibrated in 50 mM KCl, 20 mM Tris, pH 8.0. Peak fractions were pooled, dialyzed overnight at 4°C into HEK buffer (20 mM HEPES, pH 7.4, 1 mM EDTA, 50 mM KCl), aliquoted, snap-frozen in liquid N_2_, and stored at -80°C.

IQGAP1 polypeptides (6His-IQGAP1, 6His-N-IQGAP1, GST-C-IQGAP1) were expressed in *E. coli* BL21(DE3) pRARE by growing cells to log phase in TB and inducing expression with 1 mM IPTG overnight at 16°C. Cells were harvested by centrifugation and pellets stored at −80°C. Cell pellets were resuspended in lysis buffer (50 mM Potassium-phosphate, pH 8.0, 50 mM Imidazole, 500 mM KCl, 1 mM EDTA, pH 8.0, 1 mM DTT, 1% Triton X-100, 20 μg/mL DNase, lysozyme, 1 mM PMSF and a standard mixture of protease inhibitors), and kept on ice for 30 min to allow digestion, and then sonicated. Lysates were cleared for 30 min at 65,000 × *g*. For 6His-IQGAP1 and 6His-N-IQGAP1, precleared lysates were mixed with 1 ml Ni-NTA-agarose beads (Qiagen; Hilden, Germany) and incubated for 1 h rotating at 4°C. Beads were then washed three times with Ni-NTA wash buffer (20 mM Tris, pH 8.0, 50 mM Imidazole, 500 mM KCl, and 0.3% glycerol). Proteins were eluted in Ni-NTA elution buffer (Ni-NTA wash buffer plus for 500 mM Imidazole). For GST-C-IQGAP1, the precleared lysate was mixed with 1 mL glutathione-agarose beads (Thermo Fisher Scientific, Waltham, MA) and incubated for 1 h rotating at 4°C. Beads were then washed three times with GST wash buffer (20 mM Tris, pH 8.0, 500 mM KCl, and 5% glycerol) and eluted in GST elution buffer (GST wash buffer supplemented with 20 mM Reduced Glutathione (Sigma; St. Louis, MO)). All eluates were concentrated, cleared by low-speed centrifugation, and gel filtered on a Superose 12 10/300 GL column (GE Healthcare) equilibrated in HEKG_5_ buffer (20 mM HEPES, pH 7.4, 1 mM EDTA, 50 mM KCl, and 5% glycerol). Peak fractions were pooled, concentrated, snap frozen, and stored at −80°C.

SNAP-tagged IQGAP1 polypeptides (GST-SNAP-IQGAP1-6His, GST-SNAP-N-IQGAP1-6His and GST-SNAP-C-IQGAP1-6His) were purified as above for C-IQGAP1, and fluorescently labeled while still bound to the glutathione-agarose beads. For labeling, 5 μM of SNAP-surface549 or SNAP-surface649 (New England Biolabs; Ipswich, MA) was incubated with the beads rotating overnight at 4°C. The next day, beads were washed with five column volumes of GST wash buffer to remove excess dye, and then proteins were eluted with GST elution buffer. Eluates were concentrated, cleared by low-speed centrifugation, and gel filtered on a Superose 12 10/300 GL column (GE Healthcare) equilibrated in HEKG_5_ buffer. Peak fractions were pooled, concentrated, snap frozen, and stored at −80°C. For the photobleaching experiments in Figure S2B, to control for possible GST effects on the oligomerization state of full-length IQGAP1, the GST tag was removed from 549-SNAP-IQGAP1 by digestion with PreScission Protease during the labeling step above. Percent labeling of polypeptides with SNAP–surface 549 was determined by measuring fluorophore absorbance at ε560, using the extinction coefficient 140,300 M^-1^ cm^-1^. Percent labeling with SNAP–surface 649 was determined by absorbance at ε655, using the extinction coefficient 250,000 M^-1^ cm^-1^. Labeling efficiencies were consistently 55-60%.

Spectrin-actin seeds, for microfluidics-assisted TIRF, were purified from blood as described in ^29,55^. Briefly, 20 mL of packed human red blood cells (Novaseek Research, Cambridge, MA) were washed with three times with 25 mL of ice-cold Buffer A (5 mM sodium phosphate, pH 7.7, 150 mM NaCl, and 1 mM EDTA), each time centrifuging for 15 min at 2,000 × *g* at 4°C, and discarding the supernatant. To lyse cells, the cell pellet was resuspended in 700 mL (approximately 10 times the volume of washed cells) of ice-cold lysis buffer (5 mM sodium phosphate, pH 7.7 and 1 mM PMSF) and incubated for 40 min while stirring at 4°C. The lysate was centrifuged for 15 min at 45,000 × *g* at 4°C. The cloudy and viscous pellets were resuspended in wash buffer B (5 mM sodium phosphate, pH 7.7 and 0.1 mM PMSF), final volume 360 mL and homogenized by pipetting. Next, the mixture was centrifuged for 15 min at 45,000 × *g* at 4°C. The pellets were resuspended in a total volume of 180 mL of wash buffer B and homogenized as above, then centrifuged as above. This process was repeated once more. Pellets are translucent at this stage. Next, the Spectrin-actin was extracted by resuspending each pellet in 5 mL of extraction buffer (0.3 mM sodium phosphate, pH 7.6 and 0.1 mM PMSF), combining the contents into one tube, adjusting the volume to 60 mL with the same buffer, and centrifuging for 30 min at 60,000 × *g* at 4°C, repeated once. The final pellet was resuspended in an equal volume of extraction buffer and gently vortexed, then incubated for 40 min in a water bath at 37°C while manually inverting the tubes every ∼10 min. Finally, the sample was precleared for 30 min at 450,000 × *g* at 4°C. DTT (2 mM final) and protease inhibitors were added to the cleared supernatant, and an equal volume of cold glycerol (50% final concentration) was mixed into the solution. Spectrin-actin seeds were aliquoted and stored at –20°C.

### Open-flow TIRF microscopy

Glass coverslips (60 × 24 mm; Thermo Fisher Scientific) were first cleaned by sonication in detergent for 60 min, followed by successive sonications in 1 M KOH and 1 M HCl for 20 min each and in ethanol for 60 min. Coverslips were then washed extensively with H_2_O and dried in an N_2_ stream. The cleaned coverslips were coated with 2 mg/ml methoxy–poly(ethylene glycol [PEG])–silane MW 2,000 and 2 µg/ml biotin-PEG-silane MW 3,400 (Laysan Bio, Arab, AL) in 80% ethanol pH 2.0, and incubated overnight at 70°C. Flow cells were assembled by rinsing PEG coated coverslips with water, drying with N_2_, and adhering to μ-Slide VI0.1 (0.1 × 17 × 1 mm) flow chambers (Ibidi, Fitchburg, WI) with double-sided tape (2.5 cm × 2 mm × 120 µm) and 5-min epoxy resin (Devcon, Danvers, MA). Before each reaction, the flow cell was incubated for 1 min with 4 µg/ml streptavidin in HEKG_5_ buffer (20 mM Hepes pH 7.4, 1 mM EDTA, 50 mM KCl, and 5% glycerol), followed by 1 min with 1% BSA in HEKG_5_ buffer, and then equilibrated with TIRF buffer (10 mM imidazole pH 7.4, 50 mM KCl, 1 mM MgCl_2_, 1 mM EGTA, 0.2 mM ATP, 10 mM DTT, 15 mM glucose, 20 µg/ml catalase, 100 µg/ml glucose oxidase) plus 0.5% methylcellulose (4,000 cP). Finally, actin and other proteins were flowed in, as specified in figure legends.

### Microfluidics-assisted TIRF (mf-TIRF) microscopy

Actin filaments were first assembled in flow cells^19,29,42,56^. To do this, coverslips were cleaned as above (see Open-flow TIRF microscopy), and then coated with an 80% ethanol solution containing 2 mg/ml methoxy-PEG–silane MW 2,000 (adjusted to pH 2.0 with HCl) and incubated overnight at 70°C. A 40-µm-high Polydimethylsiloxane (PDMS) mold with three inlets and one outlet was mechanically clamped onto a PEG-silane–coated coverslip. The chamber was then connected to a Maesflo microfluidic flow-control system (Fluigent; Chelmsford, MA), rinsed with TIRF buffer, and incubated for 5 min with 1% BSA and 10 µg/ml streptavidin in TIRF buffer. Spectrin-actin seeds in TIRF buffer were passively absorbed to the coverslip for 10 min, then washed with TIRF buffer. Next, 1 µM G-actin (15% Alexa-488 labeled) and 5 µM profilin in TIRF buffer were introduced in order to polymerize actin filaments (with free barbed ends) from the spectrin-actin seeds. Once filaments were polymerized to a desired length (3-10 µm unless otherwise specified), specific proteins were flowed in, as described in the figure legends.

### Image acquisition and analysis

Single-wavelength time-lapse TIRF imaging was performed on a Nikon-Ti2000 inverted microscope equipped with a 150-mW Argon laser (Melles Griot), a 60× TIRF-objective with a numerical aperture of 1.49 (Nikon Instruments Inc.), and an electron multiplying charge-coupled device (EMCCD) camera (Andor Ixon; Belfast, Ireland). One pixel was equivalent to 143 × 143 nm. Focus was maintained by the Perfect Focus system (Nikon Instruments Inc.). Open-flow TIRF microscopy images were acquired every 5 s and exposed for 100 ms using imaging software Elements (Nikon Instruments Inc.; New York, NY). mf-TIRF microscopy images were exposed every 10 s (or 30 s where noted) and exposed for 100 ms using imaging software Elements (Nikon Instruments Inc.).

Images were analyzed in FIJI version 2.0.0-rc-68/1.52e (National Institutes of Health; Bethesda, MD). Background subtraction was conducted using the rolling ball background subtraction algorithm (ball radius, 5 pixels). For open-flow TIRF assays, polymerization rates were determined by plotting the filament length every 25 s and measuring the slope. For mf-TIRF assays, the depolymerization rates were determined by generating kymographs (FIJI kymograph plugin) from individual filaments. The kymograph slope was used to calculate barbed end depolymerization rates. (Rate measurements assumed one actin subunit contributes 2.7 nm to filament length). All binding curves were fit with the following hyperbolic equation (manually entered into Graphpad Prism 8.0 (San Diego, CA)):

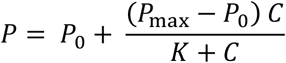

where *P* is polymerization or depolymerization rate, *P*_0_ is the rate in absence of IQGAP1 polypeptides, *P*_max_ is the rate of polymerization at saturating conditions, *K* is the IQGAP1 polypeptide concentration at half-saturation, and *C* is the IQGAP1 polypeptide concentration.

Pauses in barbed end growth were determined from traces of actin filament length versus time (Fig. 1D). A pause was defined as no change in filament length for two or more frames (5 sec per frame). The cumulative frequency of lifetime measurements of barbed end pausing was plotted and fit to a one-phase exponential association equation, used to calculate the average dwell time (Graphpad Prism 8.0).

For calculating the dwell times of GST-649-SNAP-IQGAP1, 549-SNAP-IQGAP1 (without GST tag), GST-549-SNAP-N-IQGAP1, and 549-SNAP-N-IQGAP1 molecules on actin filament sides, a kymograph was generated (using FIJI kymograph plugin) from individual sparsely decorated filaments. The lifetime measurements of the molecules were plotted, fit to a one phase exponential association equation, and used to calculate dwell times (Graphpad Prism 8.0).

For the single-molecule step-photobleaching experiments, either 2nM 649-SNAP-IQGAP1 or 2 nM 549-SNAP/Biotin-IQGAP1 in TIRF buffer without glucose oxidase and catalase was transferred into a flow cell as above, and the immobilized spots (either passively absorbed, 649-SNAP-IQGAP1, or anchored by streptavidin-biotin-PEG linkage to the slide surface, 549-SNAP/Biotin-IQGAP1) were subjected to continuous laser exposure with no delay acquisition at 100% laser power. Background fluorescence was conducted using the rolling ball background subtraction algorithm (ball radius, 5 pixels). Fluorescence intensities of individual spots were obtained by measuring the mean signal of a 6 x 6 pixel box (∼1.5 μm2) encompassing each spot. Stepwise reductions in the integrated fluorescence intensity time records of individual spots were identified and counted. The oligomeric states of GST-649-SNAP-IQGAP1 549-SNAP-IQGAP1 (without GST tag), GST-549-SNAP-N-IQGAP1, and 549-SNAP-N-IQGAP1 molecules were determined by comparing distributions of the number of photobleaching events to probability distributions for the number of fluorescent subunits calculated from the binomial distribution as described previously ^21^.

For actin filament bundling assays, 2 μM monomeric actin (10% Oregon green-labeled) was polymerized in TIRF-Buffer for 5 min at room temperature. After actin filaments were grown to 5-10 μm, IQGAP1 polypeptides were flown in and bundling was monitored for 15 min, acquiring every 5 or 10 s. Actin filament bundling was measured by subtracting background fluorescence using the rolling ball background subtraction algorithm (ball radius, 50 pixels). The segmented line tool was used to trace all actin filaments/bundles in the field of view. All line segments intensities were then normalized to the length of the measured segment (AU/μm). The intensity measurements were plotted for each time point (every 200 s). Intersections of actin filaments/bundles were excluded from segment measurements as we could not determine if the intersection was a part of the bundle. To measure the actin bundle thickness, the segmented line tool was used to draw a line perpendicular to actin filaments/bundles in the field of view. The intensity of the line segment was plotted and fit to a 2D Gaussian in FIJI. The intensity at full-width half-max (FWHM) in for each line trace was measured and recorded 1,000 s after flowing in IQGAP1 polypeptides. The thickness of the bundles in the presence of IQGAP1 polypeptides was calculated by normalizing the FWHM intensity to the control reactions.

### Quantitative Western Blotting

Western blotting was used to determine endogenous IQGAP1 proteins levels in U2OS cells (American Type Culture Collection; Manassas, VA)). Cells were pelleted and resuspended in lysis buffer (150 mM NaCl, 1.0% NP-40, 1.0% sodium deoxycholate, 1% SDS, 50 mM Tris, pH 7.5, 2 mM EDTA, 0.2 mM sodium orthovanadate, 20 mM β-glycerophosphate, 50 mM sodium fluoride, 1 mM PMSF, 1 mM DTT, and 1× Roche complete protease inhibitor mixture), and incubated at 4°C for 30 min with vortexing every 10 min. Lysates were precleared by centrifugation at 15,300 × *g* for 30 min at 4°C, and the concentration of the soluble protein fraction was determined by Bradford assay (Biorad, Hercules, CA). Known amounts of purified 6His-IQGAP1 were run on gels alongside U2OS cell lysates and blotted with a 1:1,000 dilution of rabbit anti-IQGAP1 (ab133490; Abcam, Cambridge MA). Blots were washed, probed with a 1:10,000 dilution of secondary goat anti-rabbit HRP antibody (Thermo Fisher Scientific), washed again, and then incubated for 1 min with Thermo Scientific™ SuperSignal™ West Pico PLUS Chemiluminescent Substrate (Thermo Fisher Scientific). Bands were detected on a BioRad Chemidoc MP imaging system, and quantified by densitometry using Imaging Lab version 6.0.1 software (Biorad). A standard curve for the purified protein was generated, and the amount of IQGAP1 protein in the loaded cell lysates was determined by comparison to the standard curve. Values were averaged from three independent blots. For calculations of cellular concentrations of IQGAP1, the concentration of total protein in the cytoplasm was assumed to be 100 mg/ml^51^. The amount (in grams) of IQGAP1 in 5 µg of lysate was determined, and then the molar concentration of each protein was calculated based on its known molecular weight.

